# Movement behavioral plasticity of benthic diatoms driven by optimal foraging

**DOI:** 10.1101/682153

**Authors:** Wen-Si Hu, Mingji Huang, H. P. Zhang, Feng Zhang, Wim Vyverman, Quan-Xing Liu

## Abstract

Adaptive locomotion of living organisms contributes to their competitive abilities and helps maintain their fitness in diverse environments. To date, however, our understanding of searching behavior and its ultimate cause remains poorly understood in ecology and biology. Here, we investigate motion patterns of biofilm-inhabiting marine raphid diatom *Navicula arenaria* var. *rostellata* in two-dimensional space. We report that individual *Navicula* cells display a “circular run-and-reversal” movement behavior at different concentrations of dissolved silicic acid (dSi). We show that gliding motions of cells can be predicted accurately with a universal Langevin model. Our experimental results are consistent with an optimal foraging strategy and a maximized diffusivity of the theoretical outcomes in which both circular-run and reversal behaviors are important ingredients. Our theoretical results suggest that the evolving movement behaviors of diatoms may be driven by optimization of searching behavioral strategy, and predicted behavioral parameters coincide with the experimental observations. These optimized movement behaviors are an evolutionarily stable strategy to cope with environmental complexity.

**ONE SENTENCE SUMMARY:** Novel experiments and modelling reveal that raphid diatoms can actively exploit resources in complex environments by adjusting their movement behavior.

## INTRODUCTION

The rich diversity of organisms’ movement behavior has long invoked curiosity. Plants may passively adapt their inclining positions to alleviate competition for light (1,2), while most animals and many microorganisms can actively move from one place to another to seek forage, to mate with partners (3,4), or to escape from predators (5,6). A comprehensive understanding of the drivers, patterns and mechanisms of organismal movement is central to elucidating its ecological and evolutionary significance (7–10). In the extensive body of movement behavioral ecology, particular interest has been paid to foraging, being a fundamental activity providing energy throughout an organism’s life cycle (5,8). In spite of diverse modes of foraging movement among life forms (1,8,9), their intrinsic spatiotemporal patterns may converge to maximize biological fitness of individual foragers as it is predicted by the optimal foraging theory (OFT) (11, 12).

So far, perhaps the most convincing evidence supporting this prediction is provided by theoretical and experimental studies on movement patterns of a range of microorganisms (e.g. swimming bacteria, microalgae and multi-cellular planktons) in strictly controlled microcosm environments (13–16). In these systems with low information availability, microorganisms having weak resource detection capabilities usually perform random-like movements during the process of foraging. It has been repeatedly observed that intermittent locomotion (also known as stop-and-go movement or pause-travel locomotion, Fig. 1A) is common in these cases, and is characterized by discontinuous movements interwoven with significant punctuations and reorientations. A prevalent idea is that foraging efficiency can be maximized by certain statistical properties provided by their movement patterns. For example, the probability distributions of time intervals or spatial displacements between reorientations have been found to fit Brownian type or Lévy type walks, which are theoretically regarded to be optimal solutions of the random search problems under specific conditions (13,17–23).

**Fig. 1:**
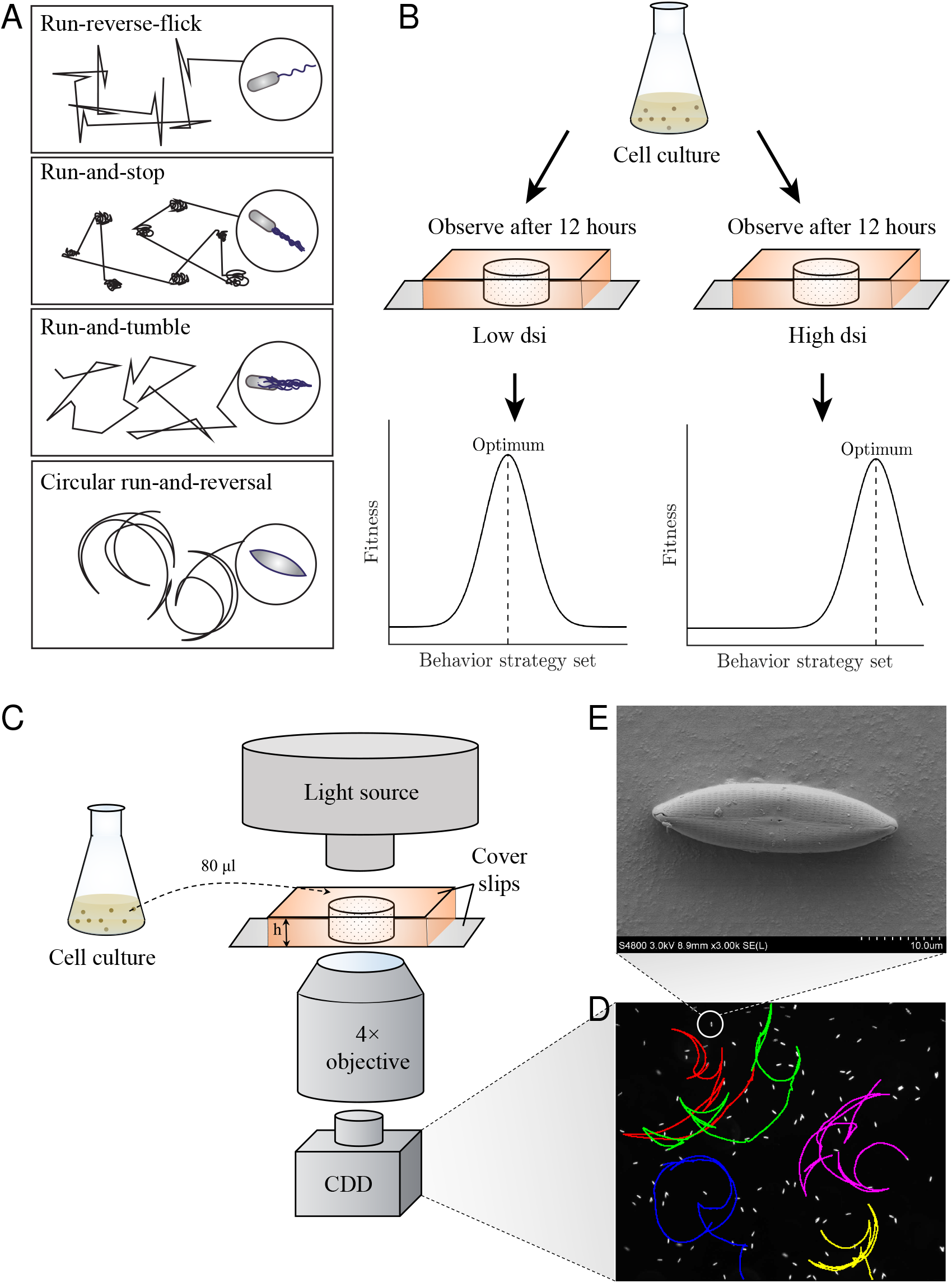
Theoretical hypothesis and experimental setup. (**A**) Three typical patterns of movement behaviors of microorganisms, showing the ‘run-and-stop’, ‘run-reverse-flick’, ‘run-and-tumble’, and the ‘circular run-and-reverse’ pattern of marine diatoms. (**B**) The characteristics of an optimization model by adjusting movement behavioral plasticity. The dash line shows peaks predicted fitness and therefore what would be expected in nature. When the environment changes, its optimal value would change accordingly. (**C**) The schematic of the experimental setup (not to scale). (**D**) An example of the observed movement trajectories. (**E**) Scanning electron microscope image of species *Navicula arenaria* var. *rostellata* shows an boat-shape cell, where the two raphes can spray the extracellular polymeric substances (EPS) to obtain self-propulsion.

The suggestion that the optimal foraging principle underpins diverse movement forms is indeed appealing. However, the universality of this strikingly simple principle remains controversial. It has been argued that there are exceptions in real-world ecosystems that are more complex deviating theoretical hypothesis (14,24,25), for instance, some individuals switching between Lévy and Brownian movement patterns as they traverse different habitat types (26,27). So far, optimal foraging is typically referred to as a hypothesis because it has not been established that the assumptions underlying these theories indeed hold. Furthermore, it remains elusive to which extent the optimal foraging principle can be verified to a range of newly discovered movement forms in microorganisms. Apart from the classical ‘run-and-tumble’ movement pattern (characterized by almost straight runs that are interrupted by tumbles) that have been extensively studied since the seminal work on *Escherichia coli* in the 1970s (10,28), recent studies discovered a variety of different movement modes (14,16,29), such as the ‘run-and-stop’ and ‘run-reverse-flick’ patterns (Fig. 1A) in the soil bacteria *Pseudomonas putida* (30), *Myxococcus xanthus* (31,32), and the marine bacteria *Vibrio alginoticus* (24). Very recently, one of the most intriguing movement patterns was found in pennate raphid diatoms, a species-rich and ecologically important group of microalgae mostly inhabiting benthic habitats in marine and freshwater environments. They move in a gliding manner, forming trajectories that highly resemble circular arcs (16). This unique movement pattern (termed as ‘circular run-and-reversal’, see Fig. 1A, results and discussion for detailed descriptions) is distinct from those of previously documented model organisms (28), whose trajectories typically consist of line segments in contrast with circular arcs. Our understanding of the statistical properties of these movement patterns is still rudimentary. In particular, it remains unknown if this type of movement conforms to optimal foraging behavior.

Here, we performed a systematic study on the novel ‘circular run-and-reversal’ behavior in the marine biofilm-inhabiting diatom *Navicula arenaria* var. *rostellata*. By combining experimental data and theoretical analyses, we demonstrate that the circular run-and-reversal behavior plays a crucial role in optimizing searching strategies. Our results suggest that in a silicon-limited environment, the diatoms can maximize their foraging efficiency by adapting the key parameters including reversal rate and rotational diffusivity and they can change the behavior strategy in a silicon-rich environment (Fig.1B).

## EXPERIMENTAL SETUP

In our experimental microcosms (Fig. 1C), enhanced motility of the diatom *Navicula arenaria* var. *rostellata* (see *Materials and Methods* for detailed descriptions on its basic information, Fig. 1E is a picture of electron microscope image of the studied species) was stimulated by exposing cells to low concentrations of dissolved silicic acid (dSi, 15 mg/L). Sample is enclosed in a sealed chamber which consists of a silicone well, and two cover slips. The well has a diameter of 0.8 cm and a height of 0.1 cm (Fig. 1C). We used a tracing technique to quantify the movement pattern of individual diatom cells (Fig. 1D and 2A). We subsequently developed a simple theoretical model that can well capture the spatial trajectories as well as the statistical properties of their foraging movement. During the experiments, cells were placed in a coverslip chamber in dSi depleted culture medium (about 15 cells/mm^2^). The movement patterns of the diatom cells were recorded by a Ti-E Nikon phase contrast microscope with high temporal-spatial resolutions (see *Materials and Methods*).

**Fig. 2:**
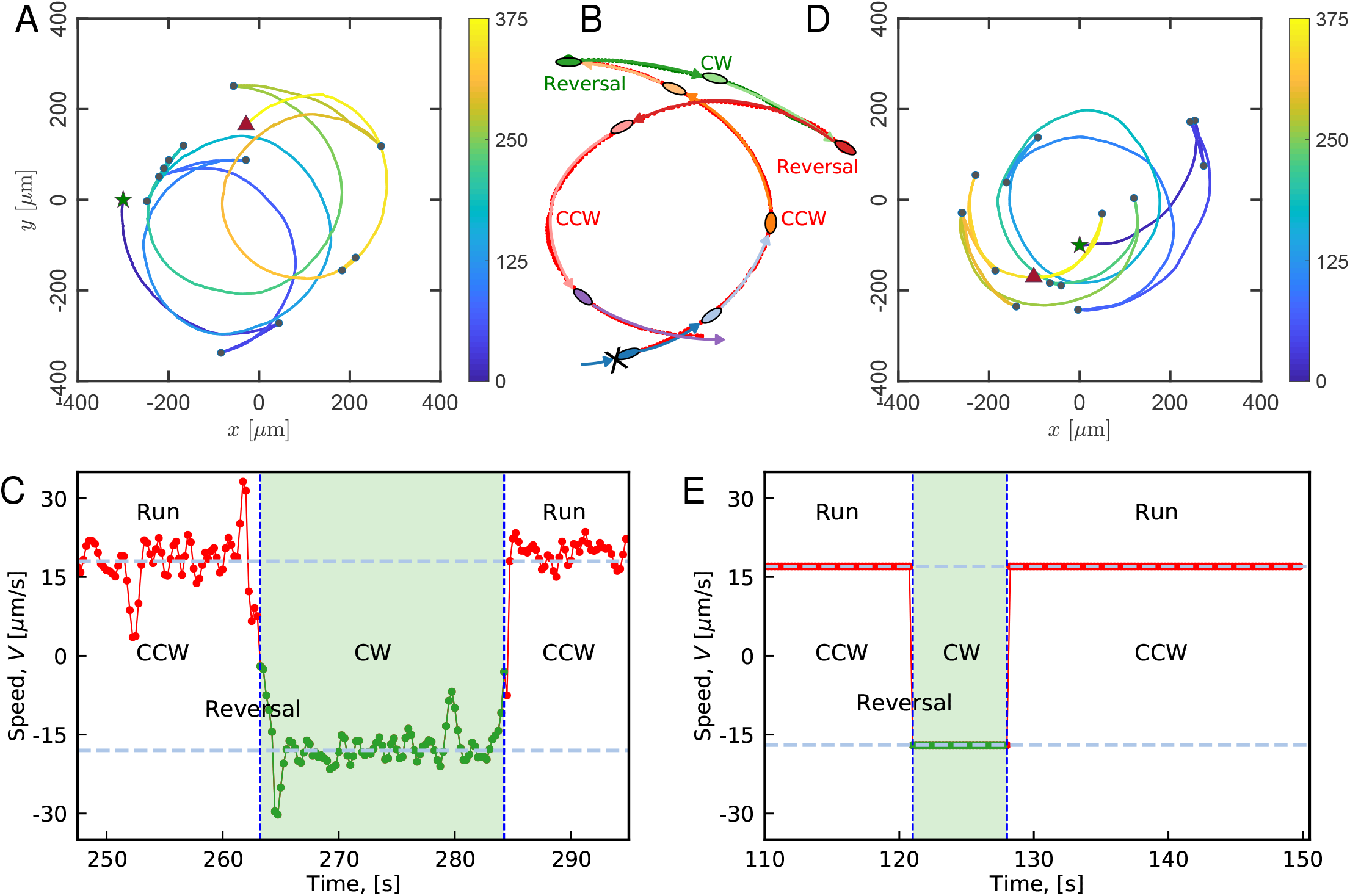
Experimental observations and theoretical predictions of the circular run-and-reversal behaviors of diatom *Navicula arenaria* var. *rostellata*. (**A**) A typical cell trajectory containing circular run and reversal behaviors captured with a microscopy at 4 frames per second (see Movie S1 for more trajectories) for 5 min. (**B**) Cropping of the partial trajectory depicts a reversal behavior with zoom in on the panel (A), where the running from CCW switches to CW through a reversal behavior, and vice versa. The arrows indicate the moving direction of the diatom cells. (**C**) Experimental data showing the movement velocity before and after a reversal occurrence; for clarity, not all speeds of the time series are shown here. (**D**) and (**E**) Predictions of spatial trajectory and reversal event obtained from model (1) with parameters value *V*_0_*=*17 μm/s, *D*_*θ*_ = 0.0054 rad^2^/s, *v* = 0.02 s^−1^, and ω = π/36 rad/s. Colorbars in panel (A, D) depict the time (see Movie S2 for theoretical simulations).

## RESULTS AND DISCUSSION

### The ‘circular run-and-reversal’ movement pattern

We observed that the movement trajectories of the diatom cells are characterized by two apparently distinguishable components: 1) continuous spatial displacements following rotation-like (resembling circular arcs) trajectories (Fig. 2A, Movie S1) in the clockwise (CW) or counter clockwise (CCW) direction; and 2) reversals of the rotational direction (Fig. 2B, Movie S1). Here we define this movement pattern as ‘circular run-and-reverse’ by adapting the term ‘run-and-tumble’ as were shown in Fig. 1A and 2.

To quantitatively characterize this ‘circular run-and-reverse’ movement pattern in a comprehensive way, we used *ca.* 30 recorded continuous individual trajectories to measure a set of key movement parameters including transitional speed, angular speed, translational diffusivity and rotational diffusivity (*D*_*θ*_, indicating the intensity of random change in particle’s orientation, which resembles the translational diffusion in space) and reversal rate (*v*, defined as the times of directional reversals per unit time). Details of the parameters are provided in Table 1.

**Table 1.**
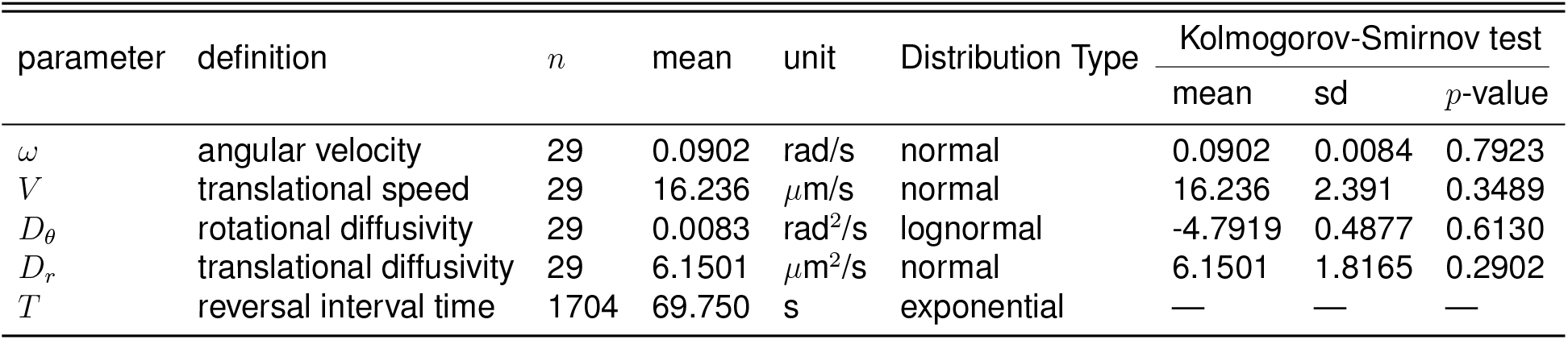
Statistical properties of measured experimental parameters on diatom movement behaviors. Experimental statistics of behavioral parameters on diatom cells at dSi concentrations of 15 mg/L. *n*, number of individuals.

In our observations, the movement speed as a function of time *V*(*t*) was around 16.2 ± 2.3 μm/s (Fig. 2C). The probability distributions of reversal time intervals of cells are well characterized by an exponential distribution with mean *T* = 1/*v* (*T* is the mean interval-reversal time, see Fig. 3A), and thus the number of reversal events in a fixed interval of time length conforms to a Poisson distribution. In addition, the statistical behavior of the rotational diffusivity (*D*_*θ*_) satisfies a Gaussian random variable with log transformation (Table 1 and Fig. S1 for the trajectories with different experimental *D*_*θ*_).

**Fig. 3:**
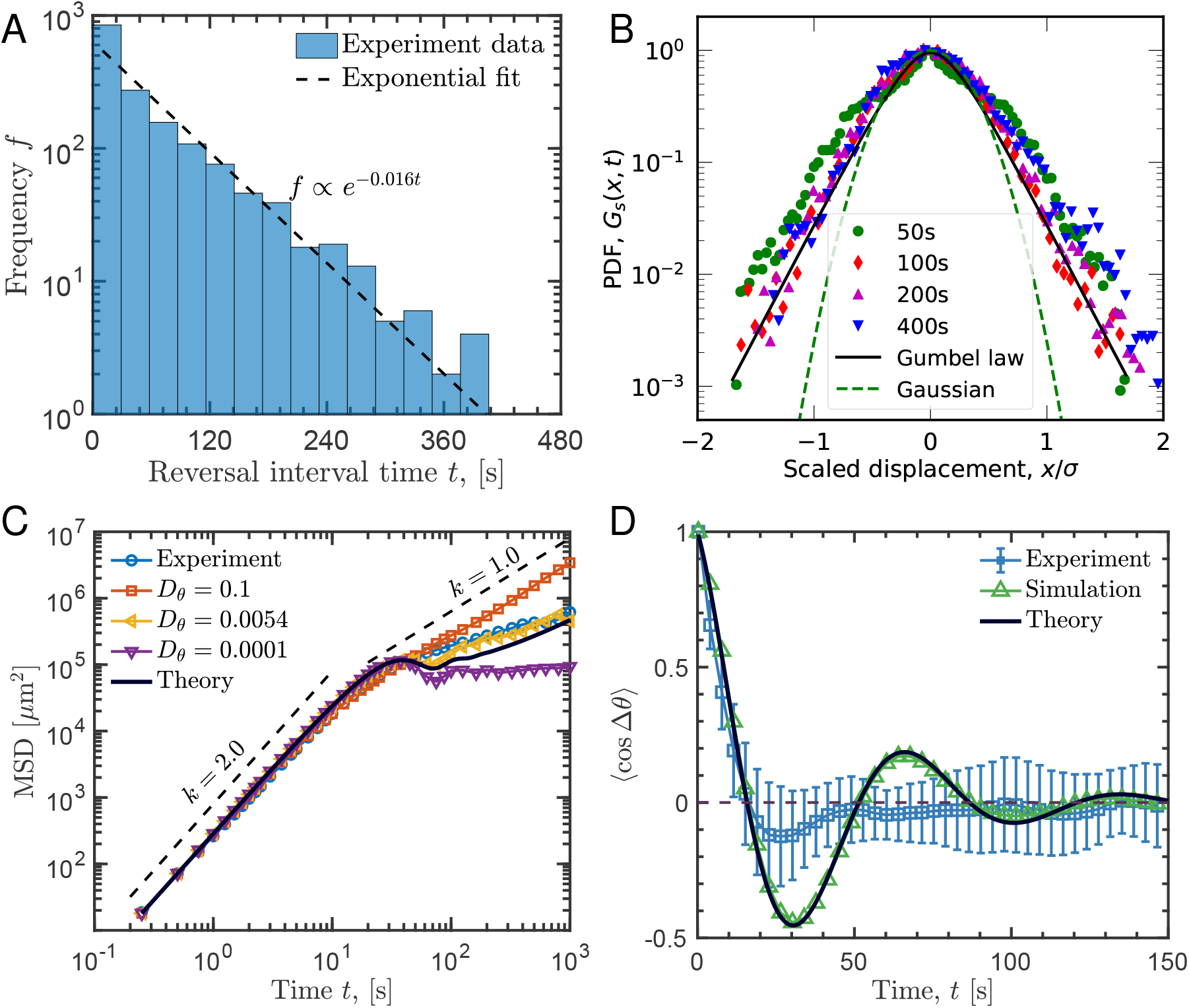
Comparing the laboratory measurements and simulation results with theoretical (analytical) predictions on diffusion behaviors of diatom cells. (**A**) Statistical distribution of 1704 reversal interval time *t* from the 29 experimental individuals trajectories, which can be well fitted by an exponential distribution *f ∝ e*^*−0.016tt*^ with the slope of −0.016. (**B**) The measured probability density functions of cells’ displacements as a function of displacement normalized by its standard deviation (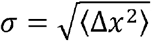) along the *x*-axis direction for different times. A fit to the data with Gumbel law (solid black lines) and Gaussian model (dashed green lines) are shown for two different time scales, where the Gumbel law of the distribution imply slower diffusion at a long-time scale. (**C**) Mean squared displacement (MSD) for three different values of the rotational diffusion coefficient *D*_*θ*_ obtained by performing the numerical simulations of model (1) and comparison with the experiments (circles symbols), respectively. By decreasing the strength of rotational diffusion in the model, the scaling behaviors of the MSD vs. time becomes consistent with confined diffusivity from ballistic behaviors similarly to cage-effect emergence after the characteristic times (~25 *s*). Parameters are *ω* = π/36 rad/s, *v* = 0.02 s^−1^, *D*_*θ*_ = 0.0054 rad^2^/s and *V*_0_ = 17 μm/s. The dashed lines are a guide to the eye to mark the change of the scaling law with 2.0 and 1.0 respectively, the solid line corresponds to the trend predicted by theory Eq. (4). (**D**) Correlation of measured and predicted changes in the direction of cells moving. Experimental data (symbols) have error bars representing lower and upper SD. Corresponding analytical predictions (solid line and dashed line with triangle symbols) are given by theory Eq. (3) and numerical simulations of model (1) respectively. The dashed line indicates 0 to guide the eye in (B).

Does the circular run-and-reverse pattern satisfy a Gaussianity? The distribution function of displacements is a fundamental statistic property for movement behavior, known as the self-part of the van Hove distribution function is defined as:

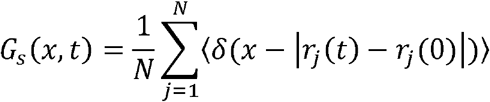

where *N* is the number of individual cells and *δ* is the Dirac delta function. They are not Gaussian behavior at long-term scales (more than 50 sec, Fig. 3B). We find that this non-Gaussian distribution can be well fitted by a Gumbel law (33):

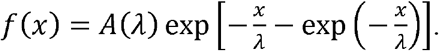

Here *λ* is a length scale, and *x* is the displacement of the cell in the *x* direction and *A*(*λ*) is a normalization constant. Therefore, we conclude that this circular run-and-reversal’ movement pattern is a non-Gaussian process for spatial searching and the rotational diffusivity leads to a subdiffusive searching behavior at long-time scales (Fig. 3C).

### Mathematical model

We developed a basic mechanistic model to capture the movement pattern of the self-propelled diatom cells at an individual level in a two-dimensional space. Adapted from the motion behavior of self-propelled non-living micro-rods (34), our discrete time model uses the abovementioned 5 movement parameters, assuming white noises with intensity *D*_*r*_ and *D*_*θ*_ in translational diffusivity and rotational diffusivity respectively. The stochastic difference equations for the movement of a single diatom cell are given by

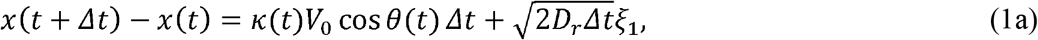

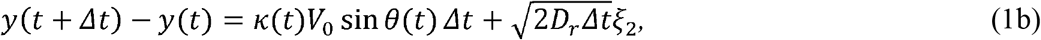

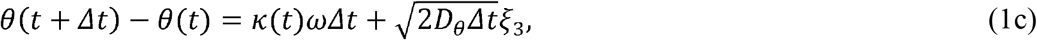

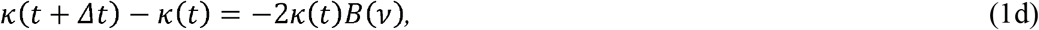

where *θ* is the direction of the movement, *x* and *y* indicate the spatial coordinate and *t* is the time. *k* is the rotational direction (*k* = 1 for CCW and −1 for CW), and ω is the angular speed. Reversal events are represented by the telegraph process, characterized by *B*(*v*) which is a Bernoulli random variable with success probability *vΔt* (*v* is reversal rate). The noise terms *ξ*_l_, *ξ*_s_ and *ξ*_3_ follow a standard normal distribution. The default values of the parameters were derived from the experimental data (Table 1).

The mathematical derivation of the effective diffusion coefficient *D* (as a measure of foraging efficiency) can be obtained by calculating the probability distribution functions *Ψ*_±_(***r***, *θ, t*) (34,35),

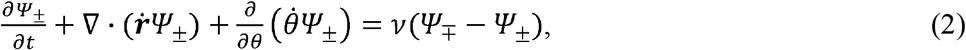

with 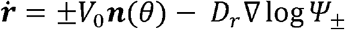, 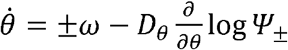. Here, the probability density function of cells’ spatial position ***r*** = (*x, y*) changes following the Fokker-Planck equation associated with the Langevin equations (Eqs. 1).

We can obtain the time-dependent expected change in orientation of the diatom cells by multiplying Eq. (2) by cos*Δθ* and separately by sin *Δθ* respectively, and then integrating both equations over *θ* and ***r***. By solving a linear system of ordinary differential equations for 〈*k* cos *Δθ*〉(*t*) and 〈*k* sin *Δθ*〉(*t*) (see Text 1 for details), the analytical prediction of temporal correlation of orientation 〈cos *Δθ*〉 is given by

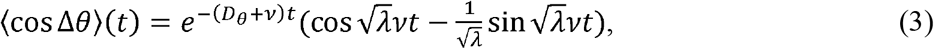

where 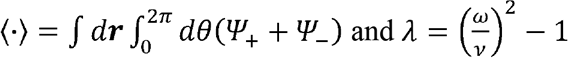.

Theoretically, we can further obtain the analytical predictions of the time-dependent mean-squared displacements (MSD) and effective diffusion coefficient from the Fokker-Planck equation (Eq. 2). Using mathematical derivation, we can obtain the analytical expression of MSD 〈**Δ*r***^2^〉(*t*) as

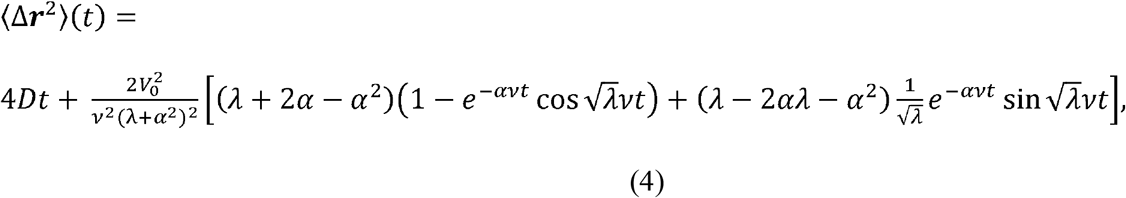

where

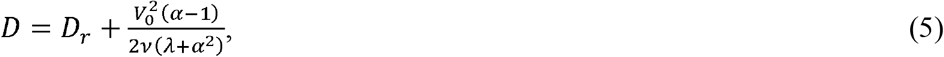

is the effective diffusion coefficient (or diffusivity for simplicity hereafter) with 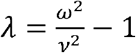 and 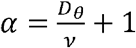. Note that for noncircular motion, i.e. *ω* = 0, *v* = 0, our model (Eq. 5) defines a system of persistent random walks characterized by a diffusivity 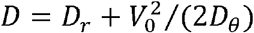 (28, 36).

### Model validation

Our model can indeed capture the spatial patterns and dynamics of the diatom movement, as reflected by the agreement between the model prediction and experiment data (Fig. 2D and E, Fig. 3C and D, movie S2). A visual check, albeit in a non-quantitative way, suggests that the circular run-and-reverse mode can be well reproduced by our model (Fig. 2D, movie S2). A rigorous validation usually requires scrutinizing essential behavioral parameters, including MSD and temporal correlation of orientation 〈cos *Δθ*〉 (34). With respect to MSD, we do find that our model (from both analytic and simulated results) is well in line with the experimental data, in the sense that they both present a highly consistent two-regime pattern of MSD as a function of time (the curves of the model results and experiment data are almost completely overlapping in Fig. 3C). At short time scales (*t* < *t*_*c*_ = 25 s), MSD as a scaling function of time shows ballistic dynamics with a scaling exponent of 2.0, whereas at long-time scales (*t* > *t*_*c*_), MSD changes to sub-diffusive behavior indicated by a scaling exponent less than 1.0. A decreasing rotational diffusivity *D*_*θ*_ leads to a decline of the scaling exponent, indicating a weakened diffusive ability when the cell motion is getting closer to the circular motion (Fig. S2). In addition, our model predicts that the diffusion behavior would converge to normal diffusion (with scaling exponent close to 1.0) after a relative long plateau even for low rotational diffusivities (see Fig. S3), which is often a general property across many diffusion behaviors. A further comparison between the model results and experiment data reveals a consistent pattern of temporal correlation of orientation 〈cos *Δθ*〉 as a function of time. Specifically, at short time scales, positive correlation coefficients are consistently found in the model and data, suggesting a positive feedback in directional persistence (Fig. 3D). The negative correlation coefficients are consistently present due to cells running a half arc associated with weak stochastic direction fluctuation. At longer time scales, correlation of orientation is dominated by noise, and the coefficients approach 0 over time (*t* < 100 *s*).

### Movement behavior driven by optimal foraging

If the prediction of the optimal foraging theory holds for the studied diatoms, one intuitive corollary is that the observed “circular run-and-reversal” movement mode can provide a statistical property that can maximize foraging efficiency. Indeed, our model analyses together with experimental data lend support to this speculation.

In the model setting with homogeneously distributed forage targets (Fig. 4A), our simulation analyses (see *Materials and Methods*) show that the amount of resource remaining in the environment decays in an exponential manner over time (Fig. 4B). We thus use the exponent of the exponential decay as a straightforward indicator of foraging efficiency, estimated by the regression slope of unconsumed resource on logarithmic scale over time. A larger exponent (*τ*) means that more resource targets can be found per unit time, hence indicating a higher foraging efficiency. However, this indicator cannot not be feasibly derived from the analytic model. Instead, we used effective diffusivity as a measure of foraging efficiency. In both analytic models and simulations, we consistently found optima of foraging efficiency at rotational diffusivity *D*_*θ*_ around 0.1 (Fig. 4C, E). A striking finding is that this optimal point is very close to the experimentally observed values of *D*_*θ*_. This is especially true for the simulations (there is only 9% deviation between the model and experiments, Fig. 4C) with a more realistic measure of foraging efficiency. The larger discrepancy between the analytical model and experimental data might be explained by the fact that the theoretically derived diffusion coefficient, although being strongly correlated, is insufficiently reflective of actual foraging efficiency under specific conditions (37).In our model, foraging efficiency does not show any peak with changing reversal rate (*v*) (Fig. 4D, F). However, the experimentally observed values of *v* are consistently located within the ranges presenting maximum foraging efficiency. The consistency between the experimental observations and model results remains robust if we plot the data and modeled foraging efficiency in two-dimensional parameter space (*D*_*θ*_, *v*), clearly showing that the experimental points fall within the optimal strategy regions (yellow regions in Fig. 5A, B).

**Fig. 4:**
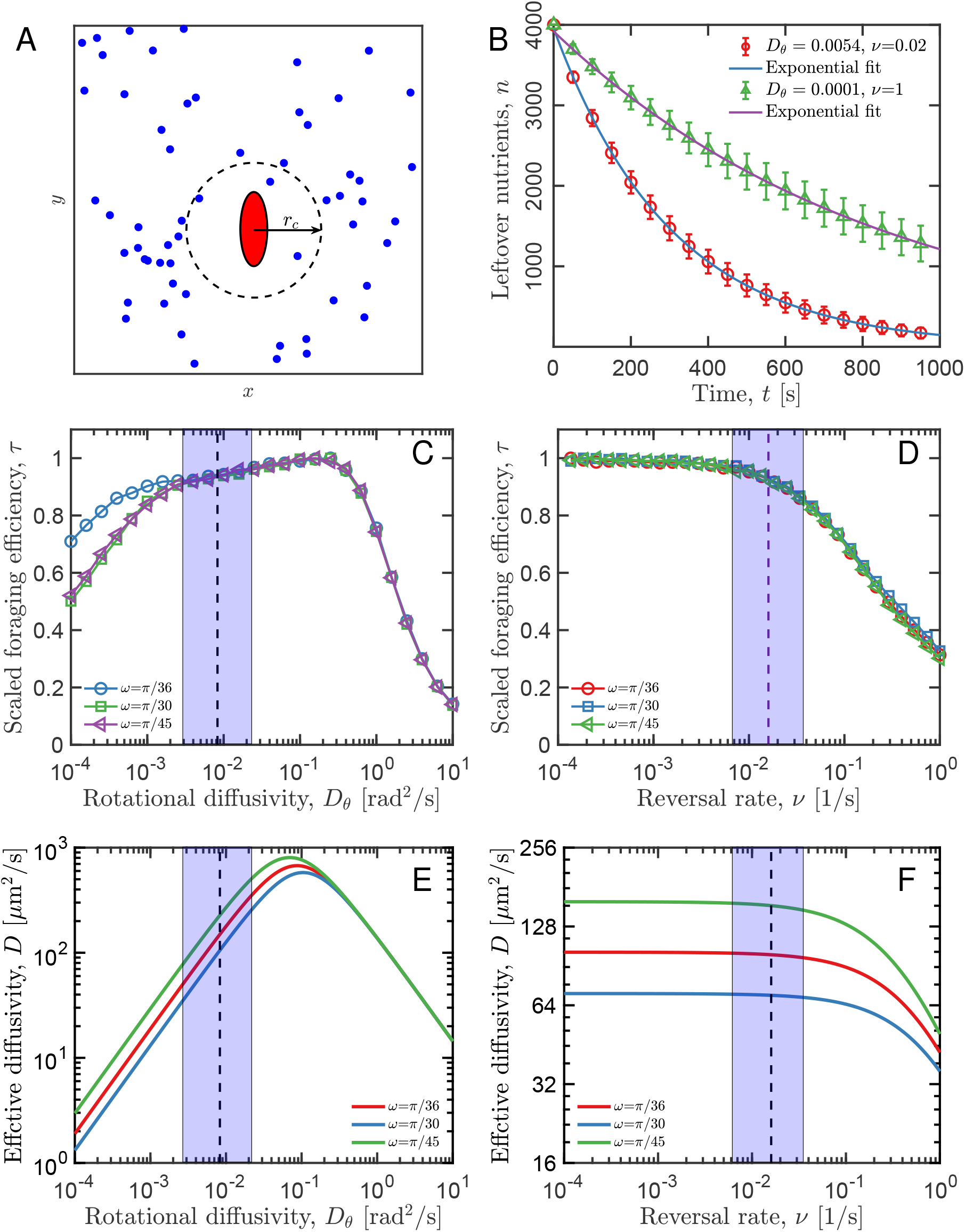
Theoretical prediction of optimal foraging strategies with spatially randomized nutrient targets. (**A**) Schematic representation (not to scale) of diatom cells blindly searching for randomly distributed nutrient resources (dots). The cells placed in a two-dimensional space move with constant speed ***V***_**0**_ and variable orientation described in model (1). The capture radius ***r***_*c*_ is about 20 ***μ***m size (dashed circle area). (**B**) The distinctive exponential function, ***n(t)*** = ***Ae***^**−*τt***^ with the decay rate *τ*, was used to describes the foraging efficiency of diatom movement strategy with respect to various value of ***D***_***θ***_ and *v*. (**C, D**) The efficiency of captured nutrients as a function of ***D***_***θ***_ and ***v***, respectively. Foraging efficiency is calculated by averaging over 1000 trajectories with various ***ω***, where the plot is scaled to the maximum value at ***D***_**θ**_ = **0.3** and ***v*** = **0.0001 s**^−1^ respectively (see fig. S6 for without scaling). (**E, F**) The analytical prediction of effective diffusivity from theory Eq. (5), coinciding with directly numerical simulations of model (1). The dashed lines and gray shaded area represent mean ±**2** SD from the experimentally measured values of ***D***_***θ***_ and ***v*** for *Navicula arenaria* var. *rostellata*.

**Fig. 5:**
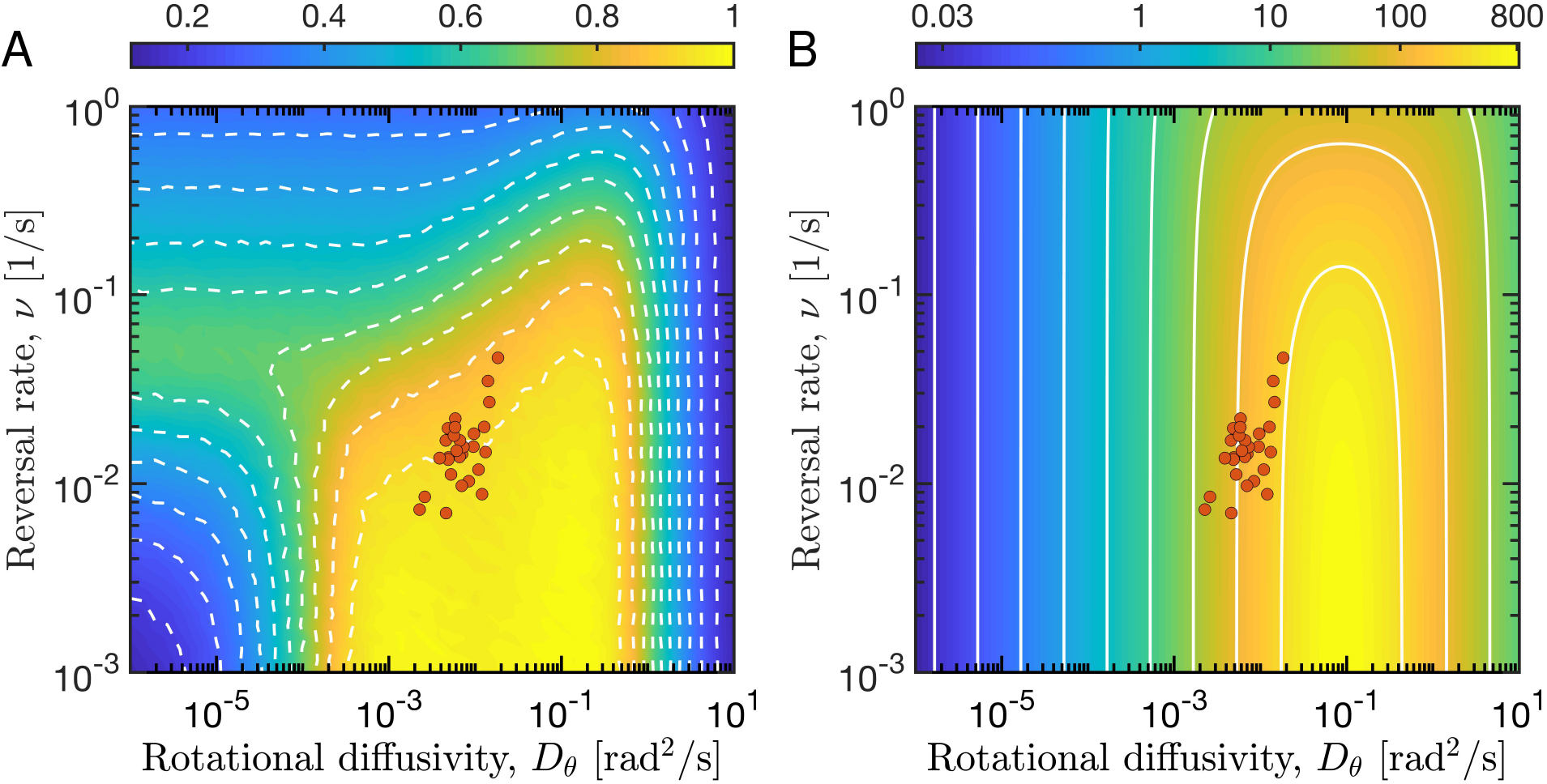
Theoretical and experimental results implicate the emergence of the foraging efficiency for various behavioral strategies. (**A**) Heatmap of foraging efficiency (colorbar) with respect to (*D*_*θ*_, *v*)-parameter space obtained from randomly distributed nutrient targets and constant movement speed for ω = π/36 rad/s and *V*_0_ = 17 μm/s. Optimal foraging occurs over a window of behavioral parameters of *v* and *D*_*θ*_, and is indicated by the yellow areas. The boundaries of the optimal regions change sharply with increasing reversal rate (white dashed lines with intervals *Δτ* = 0.1). In the low reversal rate limit, there are nonlinear effects of the rotational diffusion on diatom foraging. The colored-solid dots correspond to the experimentally measured rotational diffusion coefficients versus reversal rate on diatom *Navicula arenaria* var*. rostellata* and the colorscale indicates the scaled foraging efficiency, *τ* from 0 to 1.0. (**B**) Theoretical prediction of Eq. (**5)** on the effective diffusivity as functions of the rotational diffusivity and reversal rate. It shows a similar spatial profile comparison with directly numerical simulations.

Unlike the rotational diffusivity, the lack of optimum in foraging efficiency as a function of reversal rate *v* in our random-environment model suggests that a relatively wide range of *v* can have maximized foraging efficiency. This seems somewhat counterintuitive, and contrasts with our experimental observation presenting a rather narrow range of *v*. We infer there should be other benefit to obtain from the reversal behavior, such as self-organized biofilm formation, whereas a substantial experimental evidence is still lack.

Taken together, the experimentally observed movement parameters (both *v* and *D*_*θ*_) are consistently found in the vicinity of theoretically predicted optimal foraging efficiency. This suggests the plausibility that the movement pattern of the diatom is in line with the optimal foraging theory.

### Invasibility Analysis

If resource (forage) availability can indeed have a significant effect on the movement behavior, then the question is whether the movement strategy corresponds with an evolutionary stable strategy (ESS). We therefore generated a pairwise invasibility plot (PIP, Fig. 6), by performing an evolutionary invasibility analysis (see *Materials and Methods*) to determine whether the optimal value is a long-term outcome of competition selection, or just be exploited by free-riding strategies (38). The PIP reveals that diatom movement strategy of diatoms observed in our experiments is not only an evolutionarily stable strategy but also convergence stable. Here, we did not observe a branching to occur when the parameters of the model are changed in the evolutionary dynamics. In addition, integrating the effect of multiple attractors into evolutionary strategies remain a fascinating topic for future research. Our model provides a universal way to understand the ecologically relevant functions of movement behavior from the perspective of foraging theory.

**Fig. 6:**
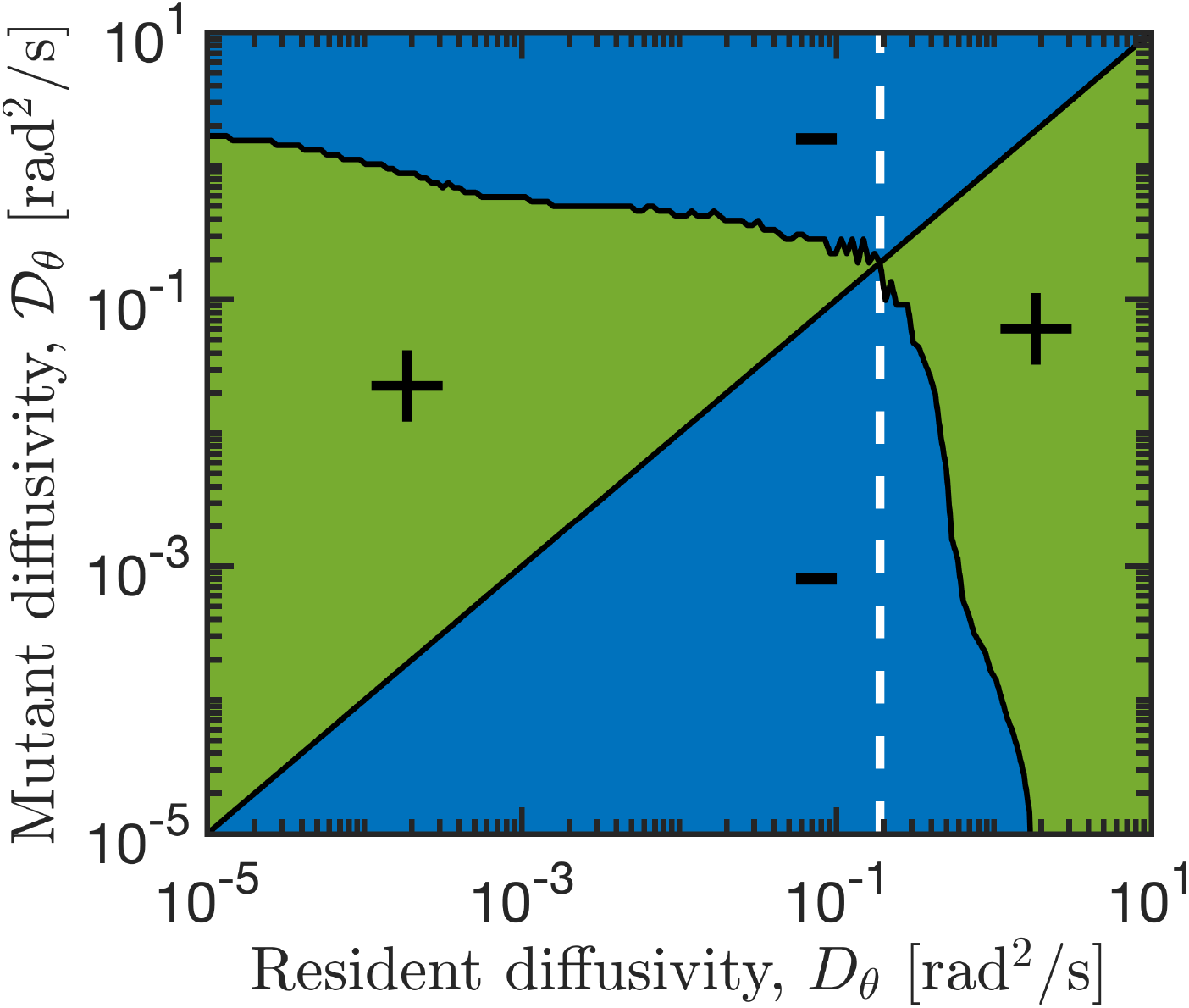
Pairwise invasibility plot (PIP) of behavioral strategy. The PIP indicates that the movement behavioral strategy of rotational diffusivity evolves toward a stable point 0.2 (vertical dashed line). For a range of resident (*x*-axis) and mutant (*y*-axis) movement strategies, the PIP describes whether a mutant has a higher (green) or a lower (blue) fitness than the resident. Plus and minus symbols indicate combinations resulting in positive and negative invasion fitness, respectively. Here, the PIP shows that the rotational diffusivity with 0.2 is the sole evolutionarily stable strategy (ESS). Simulation parameters with *v* = 0.02 s^−1^, and ω = π/36 rad/s.

## IMPLICATIONS

Our results have several useful implications. The biomechanical mechanism underlying the ‘circular run-and-reversal’ movement behavior of the diatom cells remains puzzling. A reasonable speculation is that the physical constraints of boat-shaped cells with apically located sensory receptors gliding in fluids might lead to this type of movement trajectories (39), but this is beyond the scope of this paper. Despite that, our work provides a clear demonstration that the statistical properties of this unique behavior can be ‘optimized’ towards enhanced foraging efficiency. Both theoretically and experimentally, moving beyond the statistical descriptions of movement behaviors in previous literature (13,16), our minimal model may thus serve as a useful framework for follow-up studies unravelling the ecological and evolutionary consequences of this movement behavioral plasticity in a broader context.

One fundamental question is how diatoms would adapt their movements, at individual and collective levels, in response to different foraging conditions. Indeed, our observations show that the key movement parameters revealed in our study, including reversal rate and rotational diffusivity, are sensitive to changing resource availability (see Fig. 7). The diatom cells move with low reversal rate and high effective diffusivity *D* at intermediate dSi concentrations (from 10 to 50 mg/L), whereas low and high dSi will lead to a decreased efficiency diffusivity to cells (Fig. 7A). We attribute this to the hypothesis that when silicon becomes the limiting factor, diatom cells increase searching activity to meet dSi demand for survival with a higher effective diffusivity to explore larger areas to take up dSi. It is surprising that the peak of effective diffusivity coincides with typical dSi concentrations of many coastal scenarios (Fig. 7A). The effective diffusivity shows a monotonic decline with increased reversal rates (Fig. 7B). This adaptive response suggests that diatom cells are able to sense the local dSi concentration and adjust their reversal rate to adapt to their physical surroundings. The searching efficiency within a low nutrient environment is thus strongly dependent on cell movement behaviors. Extending our results beyond dSi scavenging, there may be other attractors server as the same role to impact motion behaviors of diatoms. For instance, *in silico* comparison of experimental data led to the suggestion that diatoms have a more efficient behavioral adaptation to pheromone gradients as opposed to dSi (40). Our observations thus pave the roads for follow-up work to look further into why different movement behaviors have evolved with changing of cell body shape among diatom species, depending on cell size and shape and in response to different environmental stimuli.

**Fig. 7:**
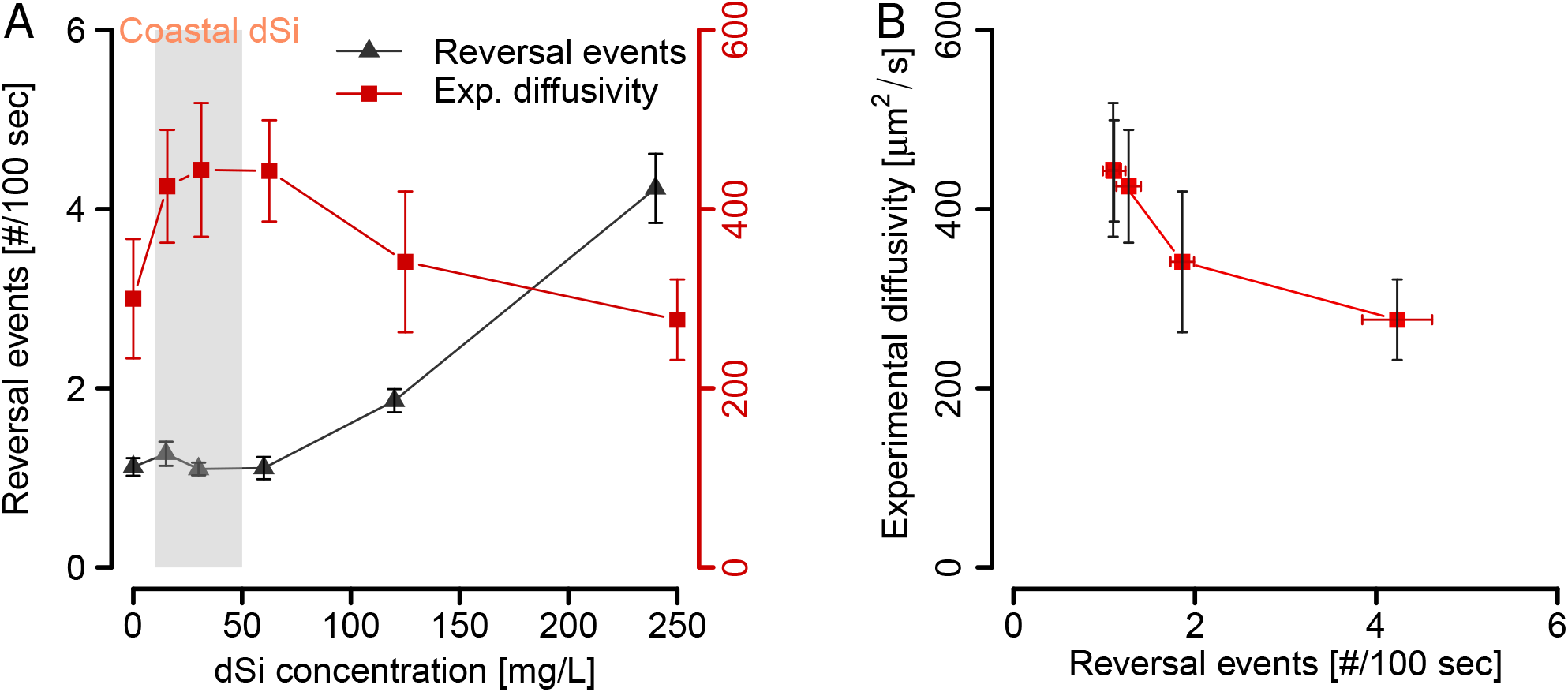
Reversal behaviors depend on the ambient dSi concentration. (**A**) The diffusivity of diatom cells maximizes at an ambient dSi concentration of about 30 mg/L and declines at low and high dSi concentrations. The reversal events show a sharply increase when dSi goes beyond 60 mg/L, but it maintains a plateau at low dSi. The grayscale rectangle indicates typical dSi concentrations in coastal ecosystems. (**B**) Efficiency diffusion coefficient, showing a monotonic decline with increased reversal events, which have a maximized dispersal coefficient about *v* = 0.02 s^−1^ coincident with model predictions.

Insights into the movement behavioral plasticity of microorganisms in aquatic environments have been generated from disciplines such as biophysics (41–43), but the focus of these studies has largely been on the statistical physical causes of behavior and not on the ultimate cause. Cases of reversal behavior were reported independently in different species of marine bacteria (24, 44, 45), and it has been suggested that it can contribute to increase foraging efficiency (24, 43) and group social effects (41), but similar evidence is still lacking for motile microalgae. This study underscores the need to study the significance of these questions in other microorganisms.

## MATERIALS AND METHODS

### Diatom cell culture and image acquisition

The *Navicula arenaria* var. *rostellata* strain 0488 (size ranges from 30~50 μm in length and 5~15 μm in width) is maintained in the BCCM/DCG diatom culture collection at Ghent University, http://bccm.belspo.be/about-us/bccm-dcg. It was isolated in January 2013 from high-nitrate intertidal flats of Paulina Schor, The Netherlands (51°21’N, 3°43’E). The isolate has since been maintained in unialgal culture in artificial seawater medium Aquil (f/2+Si). Like other naviculoid diatoms, *N. arenaria* is boat-shaped with on each valve a raphe, a specialized slit in their silica cell wall, running along its longitudinal axis. Although the precise mechanism remains unknown, diatom gliding involves an actin/myosin motility system and the secretion of adhesive EPS strands through the raphe (46).

Diatom culture were maintained using a standard protocol. One months before the experiment, cells were acclimated to 2000 Lux light intensity with a light dark cycle of 12:12 hours (INFORS HT Multitron pro, Switzerland). A 100 ml flask suspension was grown on a shaker at 20°C rotating with 100 rpm. For motility experiments, diatom cells at period of exponential phase were diluted with filtered autoclaved seawater and introduced into the test chamber for observations. The densities of individuals (about 15 cells/mm^2^) were used in order to minimize effects of cell-cell interference.

### Numerical simulations

The parameters of the simulation correspond to Fig. 2D and E, Fig. 3C. For each parameters *D*_*θ*_ and *v*, 1000 trajectories of 600 sec were simulated using a time-step *δt* = 0.1 sec with the parameter values *V*_0_ = 17 μm/s, *D*_*r*_ = 0 μm^2^/s, ω = π/36 rad/s. In Fig. 4C, *v* = 0.02 s^−1^, in Fig. 4D, *D*_*θ*_ = 0.0054 rad/s.

We start by analyzing an active cell with a sensing radius *r*_*c*_, blindly searching for 4000 nutrient resource (targets) in an environment with a homogeneous topography (47). As a diatom cruises throughout the searching space, it continuously captures nutrients that come within a capture radius *r*_*c*_ from the cells center. At each step, the ‘nutrients’ that come within a capture radius *r*_*c*_ from the cell center will be removed. We evaluate the individual search efficiency by calculating the leftover nutrients *n* in the searching space. The amount of leftover nutrients *n* in each run shows a monotonous decline as a function of the area swept by the active cell. Here, we assume that all cells use the same strategy of reversal and rotational diffusivity for the simulations. Fig. 4B plots the average amount of leftover nutrients *n*, obtained from 1,000 simulated trajectories as a function of various rotational diffusivity, so that the decay rate *τ* of the exponential fitted is defined as the foraging efficiency.

For the evolutionarily stable strategy analysis, up to 1000 cells are simulated with various prescribed rotational diffusivities *D*_*θ*_. Fitness is given by the product of survival probability and division rate. We assumed that survival probability is proportional to the foraging efficiency in the ESS analysis. A mutant strategy with a relative fitness value larger than the resident population will invade and potentially take over them. For any combination of resident and mutant movement strategy, the relative fitness of the mutants is depicted in a pairwise invasibility plot (Fig. 6).

### Calculation of time-dependent orientation correlation and MSDs of moving cells

We computed the average temporal correlation as follows: 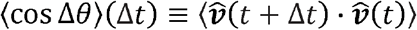, where *v̂* is the unit vector of velocity and calculated the mean square displacement via MSD(*Δt*) = 〈|***r***(*t* + *Δt*) − ***r***(*t*)|^2^〉.

## Acknowledgments

We thank Chi Xu fruitful discussion and the two anonymous reviewers for constructive comments that greatly improved the manuscript.

## Author contributions

Q.-X. L. designed research; W.-S. H. and M. H. performed research; W.-S. H. and H. P. Z. contributed new analyzed data; W.-S. H. and F. Z. contributed computer code; W.V. contributed the experimental diatom strains and experimental details; All authors contributed substantially to discussion of the content, wrote the article and reviewed and edited the manuscript before submission.

## Data and materials availability

all data needed to evaluate the conclusions in the paper are present in the paper and/or the supplementary Materials. The code to reproduce the results of this study is available on the publicly repository Dryad, DOI: 10.5061/dryad.547d7wm41.

## SUPPLEMENTARY MATERIALS

Text S1. Theoretical derivation for the expected value of MSD and diffusivity.

Fig. S1. Distribution and trajectories of experimental.

Fig. S2. Trajectories at three different values of.

Fig. S3. Behaviors of mean squared displacement (MSD) at different values of.

Fig. S4. The efficiency without scaling to the maximal foraging efficiency.

Movie S1. Typical trajectories of swimming diatoms in real experiments on diatom movement behaviors.

Movie S2. Typical trajectories of swimming diatoms in theoretical simulations of Eqs. (1).

