## Supplemental Text 1 and Fig.S1-S4 for "Movement behavioral plasticity of benthic diatoms driven by optimal foraging"

This file includes:

1. **Theoretical derivation for the expected value of MSD and diffusivity**

2. **Supplementary figures S1-S4, movies S1-S2**

Fig. S1. Distribution and trajectories of experimental $D_\theta$.

Fig. S2. Trajectories at three different values of $D_\theta$.

Fig. S3. Behaviors of mean squared displacement (MSD) at different values of $D_\theta$.

Fig. S4. The efficiency without scaling to the maximal foraging efficiency.

Movie S1. Typical trajectories of swimming diatoms in real experiments on diatom movement behaviors.

Movie S2. Typical trajectories of swimming diatoms in theoretical simulations of Eqs. (1).

### 1. Theoretical derivation for the expected value of MSD and diffusivity

We can obtain the Fokker-Planck equation from equations (1) as shown in main context, it can be given (hereafter $\boldsymbol{x} = \boldsymbol{r}$ for simplification)

$$\frac{\partial \Psi_\pm}{\partial t} + \boldsymbol{\nabla} \cdot (\dot{\boldsymbol{x}} \Psi_\pm) + \frac{\partial}{\partial \theta}\left(\dot{\theta} \Psi_\pm\right) = \nu(\Psi_\mp - \Psi_\pm), \quad \text{(S1)}$$

where

$$\dot{\boldsymbol{x}} = \pm V_0\, \boldsymbol{n}(\theta) - D_r \boldsymbol{\nabla} \log \Psi_\pm,\ \dot{\theta} = \pm\omega - D_\theta \frac{\partial}{\partial \theta} \log \Psi_\pm, \text{ here } \boldsymbol{n}(\theta) = (\cos\theta, \sin\theta).$$

Eq. S1 should hold the conditions

$\int d\boldsymbol{x} \int d\theta (\Psi_+ + \Psi_-) = 1$,

$\int d\boldsymbol{x} \int d\theta \nu(\boldsymbol{x}, \theta) \Psi_\pm \equiv \langle \nu \rangle_\pm$, and

$\int d\boldsymbol{x} \int d\theta \Psi_\pm = \langle 1 \rangle_\pm \equiv \xi_\pm$.

Multiplying each term of equation (S1) by $\cos\theta$, $\sin\theta$, $x^2$, $\boldsymbol{x} \cdot \boldsymbol{n}$, $\boldsymbol{x} \cdot \frac{d\boldsymbol{n}}{d\theta}$ respectively, and integrating, we obtain the following respectively,

$$\frac{d}{dt}\langle \cos\theta \rangle_\pm \pm \omega \langle \sin\theta \rangle_\pm + D_\theta \langle \cos\theta \rangle_\pm = \nu(\langle \cos\theta \rangle_\mp - \langle \cos\theta \rangle_\pm), \quad \text{(S2)}$$

$$\frac{d}{dt}\langle \sin\theta \rangle_\pm \mp \omega \langle \cos\theta \rangle_\pm + D_\theta \langle \sin\theta \rangle_\pm = \nu(\langle \sin\theta \rangle_\mp - \langle \sin\theta \rangle_\pm), \quad \text{(S3)}$$

$$\frac{d}{dt}\langle x^2 \rangle_\pm \mp 2V_0 \langle \boldsymbol{x} \cdot \boldsymbol{n} \rangle_\pm - 4D_r \xi_\pm = \nu(\langle x^2 \rangle_\mp - \langle x^2 \rangle_\pm), \quad \text{(S4)}$$

$$\frac{d}{dt}\langle \boldsymbol{x} \cdot \boldsymbol{n} \rangle_\pm \mp V_0 \xi_\pm \mp \omega \left\langle \boldsymbol{x} \cdot \frac{d\boldsymbol{n}}{d\theta} \right\rangle_\pm + D_\theta \langle \boldsymbol{x} \cdot \boldsymbol{n} \rangle_\pm = \nu(\langle \boldsymbol{x} \cdot \boldsymbol{n} \rangle_\mp - \langle \boldsymbol{x} \cdot \boldsymbol{n} \rangle_\pm), \quad \text{(S5)}$$

$$\frac{d}{dt}\left\langle \boldsymbol{x} \cdot \frac{d\boldsymbol{n}}{d\theta} \right\rangle_\pm \pm \omega \langle \boldsymbol{x} \cdot \boldsymbol{n} \rangle_\pm + D_\theta \left\langle \boldsymbol{x} \cdot \frac{d\boldsymbol{n}}{d\theta} \right\rangle_\pm = \nu\left(\left\langle \boldsymbol{x} \cdot \frac{d\boldsymbol{n}}{d\theta} \right\rangle_\mp - \left\langle \boldsymbol{x} \cdot \frac{d\boldsymbol{n}}{d\theta} \right\rangle_\pm\right). \quad \text{(S6)}$$

When we define the initial state as

$\Psi_+(\theta, \boldsymbol{x}, t = 0) = \delta(\theta)\delta(\boldsymbol{x})$, and

$\Psi_-(\theta, \boldsymbol{x}, t = 0) = 0$,

then, we can get the expectation of velocity-correlation

$$\int d\boldsymbol{x}_1 \int d\theta_1 \int d\boldsymbol{x}_2 \int d\theta_2 \big(V_0\, \boldsymbol{n}(\theta_1)\, \Psi_+(\theta_1, \boldsymbol{x}_1, t=0) - V_0\, \boldsymbol{n}(\theta_1)\, \Psi_-(\theta_1, \boldsymbol{x}_1, t=0)\big) \cdot$$
$$\big(V_0\, \boldsymbol{n}(\theta_2)\, \Psi_+(\theta_2, \boldsymbol{x}_2, t) - V_0\, \boldsymbol{n}(\theta_2)\, \Psi_-(\theta_2, \boldsymbol{x}_2, t)\big)$$

$$= d\boldsymbol{x}_2 \int d\theta_2 V_0^2 \cos \int \theta_2 \big(\Psi_+(\theta_2, \boldsymbol{x}_2, t) - \Psi_-(\theta_2, \boldsymbol{x}_2, t)\big) = V_0^2 (\langle\cos\theta\rangle_+ - \langle\cos\theta\rangle_-)$$

$$\equiv V_0^2 \langle\cos\theta\rangle. \tag{S7}$$

Then we get the normalize the velocity-correlation as

$$\langle\cos\theta\rangle \equiv \langle\cos\theta\rangle_+ - \langle\cos\theta\rangle_-. \tag{S8}$$

If we write the relation

$$\langle\sin\theta\rangle \equiv \langle\sin\theta\rangle_+ + \langle\sin\theta\rangle_-, \tag{S9}$$

Thus we can obtain the as follows equations

$$\frac{d}{dt}\langle\cos\theta\rangle + \omega\langle\sin\theta\rangle + D_\theta\langle\cos\theta\rangle = -2\nu\langle\cos\theta\rangle, \tag{S10}$$

$$\frac{d}{dt}\langle\sin\theta\rangle - \omega\langle\cos\theta\rangle + D_\theta\langle\sin\theta\rangle = 0, \tag{S11}$$

with the initial conditions

$t = 0,\ \langle\cos\theta\rangle = 1$, and $\langle\sin\theta\rangle = 0$.

Solving the Eq. S10 and S11, we can get

$$\langle\cos\theta\rangle(t) = e^{-(D_\theta+\nu)t}(\cos\sqrt{\lambda}\nu t - \frac{1}{\sqrt{\lambda}}\sin\sqrt{\lambda}\nu t), \tag{S12}$$

with

$$\lambda = \frac{\omega^2}{\nu^2} - 1.$$

From

$$\langle x^2\rangle \equiv \langle x^2\rangle_+ + \langle x^2\rangle_-, \tag{S13}$$

$$\langle \boldsymbol{x}\cdot\boldsymbol{n}\rangle \equiv \langle \boldsymbol{x}\cdot\boldsymbol{n}\rangle_+ - \langle \boldsymbol{x}\cdot\boldsymbol{n}\rangle_-, \tag{S14}$$

$$\left\langle \boldsymbol{x}\cdot\frac{d\boldsymbol{n}}{d\theta}\right\rangle \equiv \left\langle \boldsymbol{x}\cdot\frac{d\boldsymbol{n}}{d\theta}\right\rangle_+ + \left\langle \boldsymbol{x}\cdot\frac{d\boldsymbol{n}}{d\theta}\right\rangle_- \;, \tag{S15}$$

we can obtain the equations

$$\frac{d}{dt}\langle x^2\rangle - 2V_0\langle \boldsymbol{x}\cdot\boldsymbol{n}\rangle - 4D_r = 0, \tag{S16}$$

$$\frac{d}{dt}\langle \boldsymbol{x}\cdot\boldsymbol{n}\rangle - V_0 - \omega\left\langle \boldsymbol{x}\cdot\frac{d\boldsymbol{n}}{d\theta}\right\rangle + D_\theta\langle \boldsymbol{x}\cdot\boldsymbol{n}\rangle = -2\nu\langle \boldsymbol{x}\cdot\boldsymbol{n}\rangle, \tag{S17}$$

$$\frac{d}{dt}\left\langle \boldsymbol{x}\cdot\frac{d\boldsymbol{n}}{d\theta}\right\rangle + \omega\langle \boldsymbol{x}\cdot\boldsymbol{n}\rangle + D_\theta\left\langle \boldsymbol{x}\cdot\frac{d\boldsymbol{n}}{d\theta}\right\rangle = 0, \tag{S18}$$

with the initial conditions

$t = 0,\ \langle x^2\rangle = 0,\ \langle \boldsymbol{x}\cdot\boldsymbol{n}\rangle = 0$, and $\left\langle \boldsymbol{x}\cdot\frac{d\boldsymbol{n}}{d\theta}\right\rangle = 0$.

Solving the S16, S17, and S18, we can get

$$\langle x^2\rangle(t) = 4Dt + \frac{2V_0^2}{\nu^2(\lambda+\alpha^2)^2}\Big[(\lambda + 2\alpha - \alpha^2)\big(1 - e^{-\alpha\nu t}\cos\sqrt{\lambda}\nu t\big) + (\lambda - 2\alpha\lambda - \alpha^2)\frac{1}{\sqrt{\lambda}}e^{-\alpha\nu t}\sin\sqrt{\lambda}\nu t\Big], \tag{S19}$$

where

$$D = D_r + \frac{V_0^2(\alpha-1)}{2\nu(\lambda+\alpha^2)}, \tag{S20}$$

and $\lambda = \frac{\omega^2}{\nu^2} - 1, \alpha = \frac{D_\theta}{\nu} + 1$.

Now, we have obtained the theoretical expression of the effective diffusivity Eq. (S20).

#### 2. Supplementary figures and movies

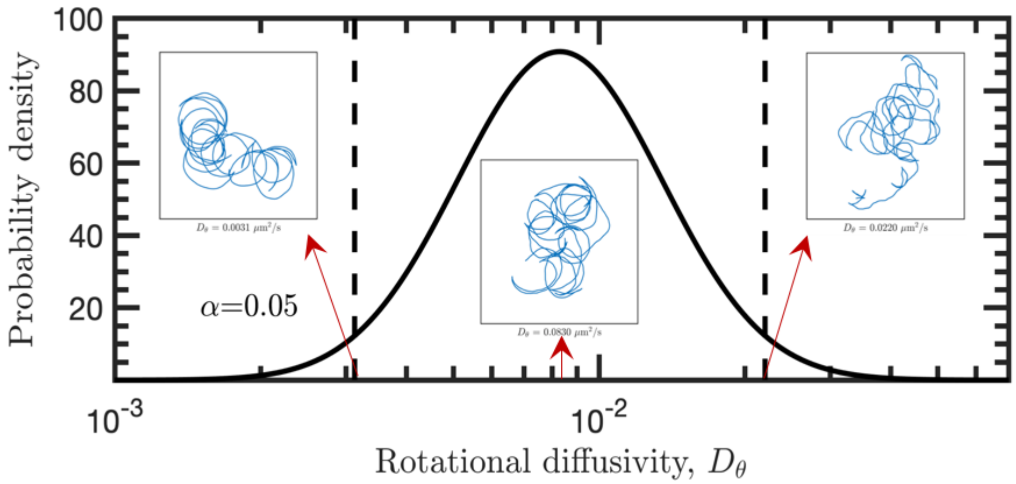

**Figure S1. Distribution and trajectories of experimental $D_\theta$.** The fitting curve of rotational diffusivity $D_\theta$ with 29 individuals which obeys log-normal distribution with 95% confidence interval. The dash lines are mean $\pm 2$SD. Inset panels show the typical trajectories at $D_\theta = 0.0031, \ 0.0830, 0.0220\ \mu m^2/s$, respectively.

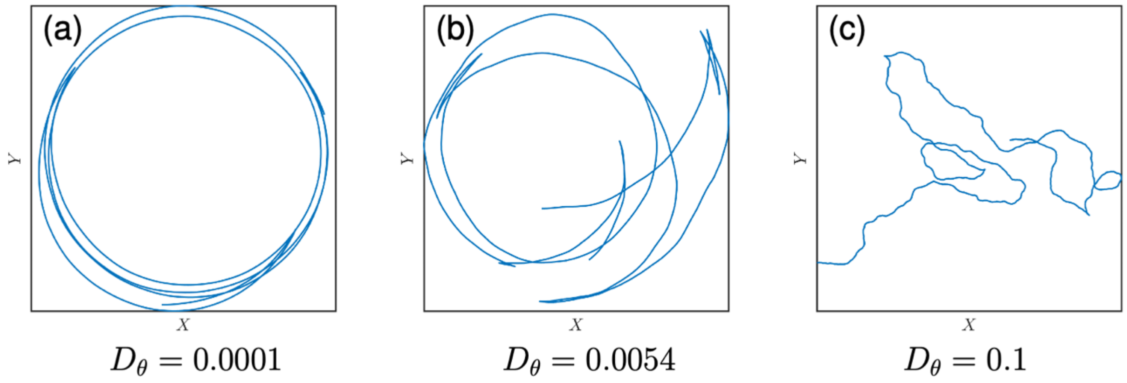


**Figure S2. Trajectories at three different values of $D_\theta$.** Trajectories at three different values of the rotational diffusion coefficient $D_\theta$. (a)-(c) depict the specific trajectories result from Eqs. (1) with the parameter $V_0$=17 $\mu m$, $\omega = 36/\pi$, $D_r = 0$ and $\nu$= 0.02 during 375 seconds. The trajectory changes from the circular motion behavior to the Brownian-like motion when $D_\theta$ increases.

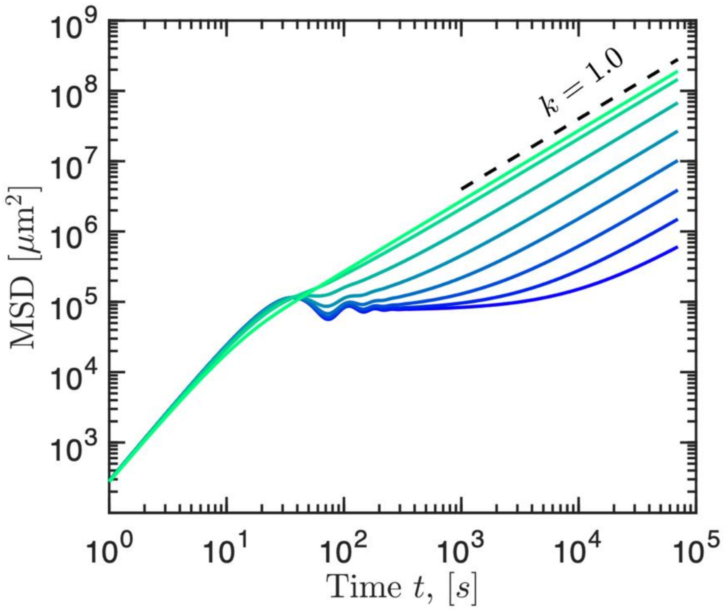

**Figure S3**. **Behaviors of mean squared displacement (MSD) at different values of $\boldsymbol{D_\theta}$.** Behaviors of mean squared displacement (MSD) at different values of rotational diffusion coefficient $D_\theta$ varies from 0.0001 to 0.1 $\mu m^2/s$ of Eq. (4). The scaling behavior shifts to normal diffusion from plateau behavior at long-term scale, called cage effect at low $D_\theta$. Diatom exhibit nearly ballistic behavior over short times and sub-diffusive behavior over time longer than about 25 second. In the last, all of them become the normal diffusive behavior.

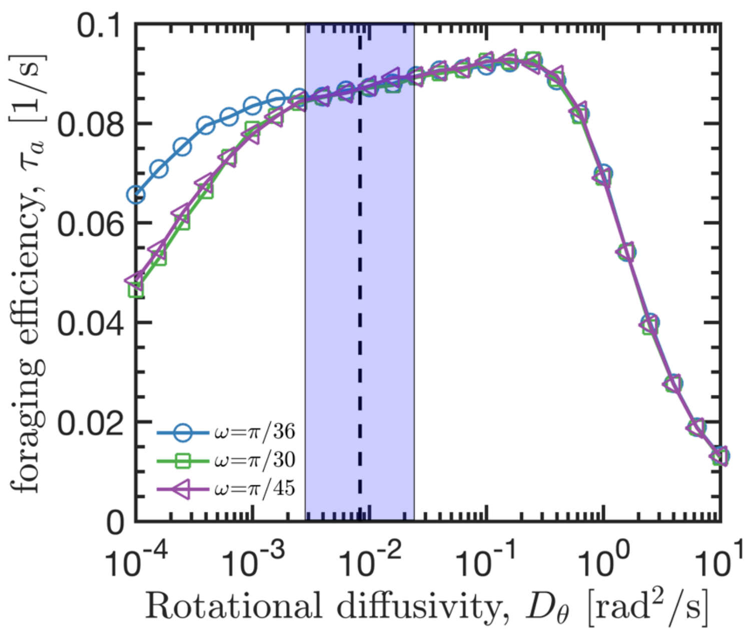


**Figure S4. The efficiency without scaling to the maximal foraging efficiency.** The efficiency without scaling to the maximal foraging efficiency of captured nutrients as a function of $\boldsymbol{D_\theta}$, respectively. The efficiency values without are averaged over 1000 trajectories at stable capture rates with various $\boldsymbol{\omega}$, where the plot is scaled to the maximum value at $\boldsymbol{D_\theta = 0.3}\mu m^2/s$ . The grey shaded area represents mean $\pm 2$SD around the experimentally measured values of $\boldsymbol{D_\theta}$ and $\boldsymbol{\nu}$ on species *Navicula arenaria* var. *rostellata*, respectively.

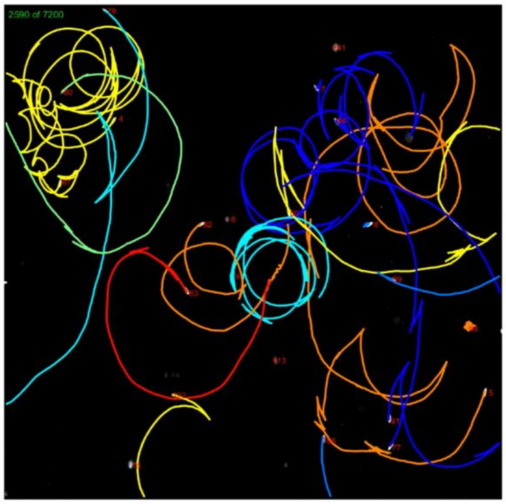


**Movies S1: Typical trajectories of swimming diatoms in experiments.** The video is played 7.25 times faster (the real recording interval is 0.25s between continued frames). The cells are marked with colored labels for a schematic visibility.

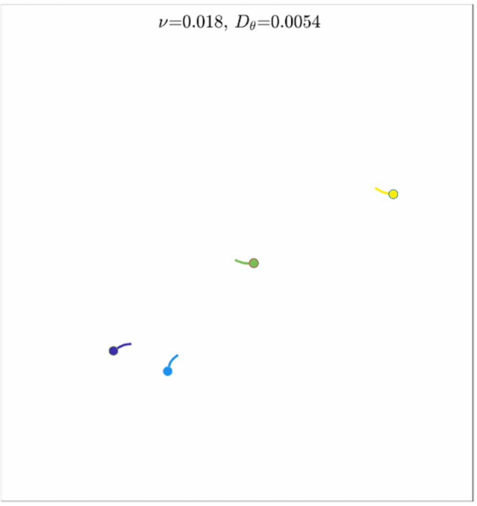


**Movies S2: Typical trajectories of swimming diatoms in simulations of Eqs. (1) in the main text.** The video is played 7.25 times faster (the real recording interval is 0.25s between continued frames). The cells are marked with colored labels for a schematic visibility.
